# Membrane contacts between caveolae and the endoplasmic reticulum regulate uptake and metabolic trapping of long-chain fatty acids

**DOI:** 10.64898/2025.12.22.695939

**Authors:** Sebastian Rönfeldt, Elin Larsson, Björn Morén, Alexander J. Craig, Lindon W. K. Moodie, Robert G. Parton, Richard Lundmark

## Abstract

The vital process of cellular fatty acid metabolism involves uptake, metabolic trapping and storage of long-chain fatty acids (LCFAs) in different lipid species. Although several different enzymes like transporters and acyl-CoA synthetases are involved, the sequential mechanistic steps of the process are poorly understood. Here, we have addressed the role of the caveola coat protein caveolin1 and the acyl-CoA synthetase FATP1 in the process as both have major impacts in the uptake and storage of fatty acids *in vivo* and *in vitro*. Using live cell microscopy and mass spectrometry-based quantification of LCFA metabolism, we found that FATP1-mediated uptake and metabolic processing of LCFAs is coupled to, and dependent on caveolin1. Furthermore, both proteins stimulate the incorporation of LCFA into phospholipids and triacylglycerol, but not in sphingolipids. Using correlative light and electron microscopy we found that membrane contact sites are formed between FATP1-enriched endoplasmic reticulum (ER) and caveolae. Analysis of their temporal stability showed that they are dynamic but frequently persist over minutes. We propose that caveolae directly couple the plasma membrane (PM) to the ER for directed LCFA transport and metabolic processing.

## Introduction

Uptake and metabolic processing of long-chain fatty acids (LCFAs) is fundamental to cell biology, facilitating ATP generation, membrane lipid synthesis and triacylglycerol (TAG) storage. Malfunctions in the control of these processes are directly linked to diseases such as obesity, diabetes and cancer (Kim & Scherer, 2021). LCFA uptake involves several mechanistic steps. First, LCFAs are released from serum albumin or from lipoprotein particles by lipases and transported across the plasma membrane (PM). This process is mediated via concentration-based membrane incorporation and lipid flipping, as well as protein-mediated transport, leading to LCFA molecules localizing in the cytoplasmic leaflet of the PM (Glatz *et al*, 2010). This allows acyl-CoA synthetases (ACSs) to metabolically trap LCFAs by incorporating coenzyme A (CoA) via thioesterification. The resulting acyl-CoA conjugates are less likely to flip/flop or to be transported out of the cell (Fig 1A). They can then be used for ATP generation in mitochondrial β-oxidation or be incorporated into monoacylglycerol (MAG), diacylglycerol (DAG) in the endoplasmic reticulum (ER). DAG can be further processed to TAG by diacylglycerol O-acyltransferase 2 (DGAT2) or to glycerophospholipids by the addition of a head group. Acyl-CoA can also be incorporated into sphingosine by ceramide synthases to generate sphingolipids in the ER (Fig 1A) (Ahmadian *et al*, 2007).

**Figure 1.**
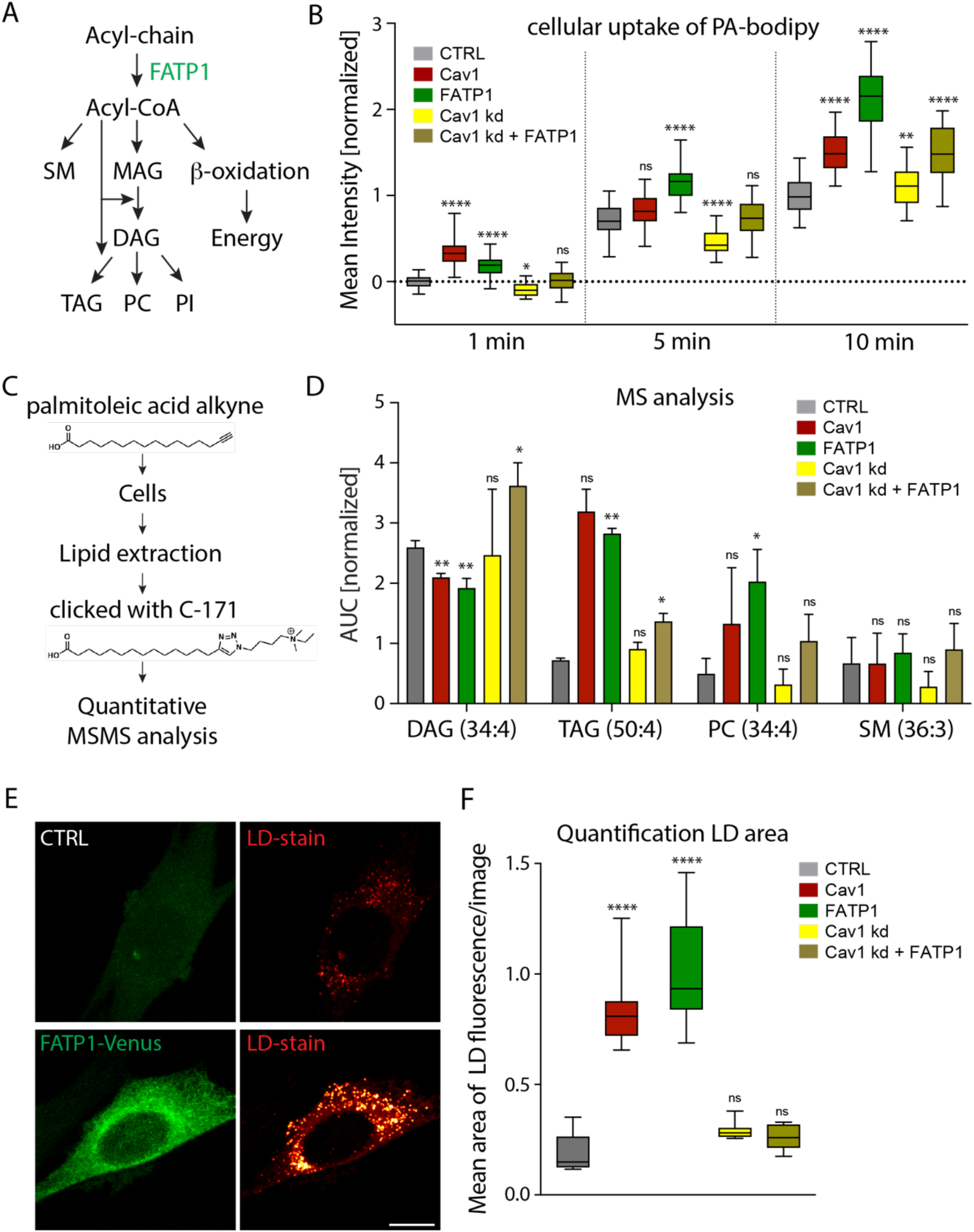
Increased uptake and metabolic processing of palmitoleic acid mediated by FATP1 is dependent on Cav1. **(A)** Schematic illustration of the metabolic processing of acyl-chains including β-oxidation in mitochondria and metabolic conversion into the derivatives Acyl-CoA, monoacylglycerol (MAG), diacylglycerol (DAG), triacylglycerol (TAG), sphingomyelin (SM), phophatidylcholin (PC) and phosphatidylinositol (PI). Metabolic enzymes investigated in this study are indicated in green. **(B)** Uptake of PA-Bodipy in wt (grey) 3T3-L1 and 3T3-L1 cells transfected with Cav1-mRFP (red) or FATP1-Venus (green), subjected to Cav1 knockdown (kd, yellow), or rescued with FATP1-Venus transfection after Cav1 kd (brown). Fluorescence mean intensity was measured at 1, 5 and 10 min, normalized to control. **(C)** Schematic illustration of palmitoleic acid alkyne (PA-alk) and the sequential steps of addition to 3T3-L1 cells, lipid extraction, chemical coupling to the C-171 click-MS reporter and analysis of metabolic derivates of the PA-alk by quantitative mass spectrometry (MS) with signature fragmentation in tandem MS (MSMS). **(D)** Bar plot of normalized area under the curve (AUC) of clicked lipid species from 3T3-L1 cells after 10 min treatment with PA-alk identified by MS analysis. The AUC from integrated extracted ion chromatograms of depicted lipid species was calculated using MassProfiler and was normalized to the AUC of the internal standard Ceramide 16:0 d30. Acyl chain length and double bonds in brackets. The bars represent the mean from three replicas ± SD. **(E)** Representative epifluorescence micrographs of control 3T3-L1 cell and a cell expressing FATP1-Venus as indicated and stained for lipid droplets (LD) using Bodipy 493/503 following 18 hours of incubation with oleic acid coupled to BSA. Scale bar 10 µm. **(F)** Quantification of the thresholded LD fluorescence area per 2.7 x 2.7 mm image (for examples see Fig S2F) of cells overexpressing or underexpressing proteins as indicated and treated with oleic acid for 18 h. Statistical significance in A and F: *, p < 0.05; **, p < 0.01, ***, p < 0.001, ****, p < 0.0001, ns, non-significant (p > 0.05).

PM transport and metabolic trapping by CoA esterification are key rate limiting steps for LCFA uptake (Glatz *et al*., 2010). Yet, the molecular and spatial details of how the hydrophobic FAs traverse the aqueous environment between the PM and ER or mitochondria remains elusive. Multiple proteins including CD36, caveolin-1 (Cav1) and the fatty acid transport proteins (FATPs) FATP1 and FATP4 have been proposed to act as LCFA transporters (Glatz *et al*., 2010). However, the mechanistic understanding of how any of these proteins facilitate fatty acid import is lacking. LCFAs readily flip between the two PM leaflets (Kamp *et al*, 2003), suggesting that proteins that facilitate inner leaflet enrichment as well as acyl-CoA incorporation promote LCFA uptake. Indeed, both vectorial acylation (Mashek & Coleman, 2006; Zhan et al, 2012) as well as protein-mediated PM transport (Glatz et al., 2010), have been proposed to drive the process and these mechanisms likely work in concert. Notably, it remains unclear whether acylation occurs directly at the PM, at internal membranes or at both sites.

The six human FATP homologues (also known as very long-chain acyl-CoA synthetases, ACSVLs) belong to the ACS superfamily based on sequence homology (Ellis *et al*, 2010; Watkins, 2008). FATPs are membrane-anchored proteins that facilitate FA adenylation using ATP, followed by thioesterification leading to CoA incorporation. However, their ACS activity is lower in comparison with other members of the family, suggesting the need for coactivity (Glatz *et al*., 2010). FATPs are differentially expressed between tissue types and localize to separate internal membranes in cells, suggesting that they channel LCFAs to different metabolic fates (Anderson & Stahl, 2013). This raises the question of how LCFAs are transported to the ER or mitochondria for acylation by FATPs when they have been incorporated into the inner PM leaflet. Cytosolic fatty acid binding proteins (FABPs) can bind and shuttle LCFAs from the PM (Glatz *et al*., 2010). However, this way of transport is not efficient enough to be solely responsible for LCFA transport and the feedback control of such a system is problematic. Therefore, it has been proposed that direct membrane contact sites (MCSs) between the PM and the ER serve as a major route of LCFA transport (Fullekrug *et al*, 2012).

Adipocytes are specialized to serve as the main energy reserve in vertebrates by metabolizing LCFAs into TAGs for storage in lipid droplets (LDs) (Ahmadian *et al*., 2007). During adipocyte differentiation, enzymes involved in LCFA metabolism are majorly upregulated. This includes the acyl-CoA synthetase FATP1 (also called ACSVL4/SLC27A1) and Cav1, which is one of the most abundant integral membrane proteins in adipocytes. Cav1, together with the peripheral protein cavin1, make up the protein foundation of the small (50 nm) PM invaginations called caveolae, which can constitute up to 50 % of the adipocyte cell surface (Parton, 2018). Apart from being a caveola component, Cav1 has also been detected in the PM outside of caveolae as well as in LDs. Collective evidence shows that both FATP1 and Cav1 play important roles in fatty acid (FA) metabolism in adipocytes (Pilch et al. 2011). Additionally, insulin-dependent FA uptake is blocked in FATP1 knock-out (KO) mice and these animals are resistant to high fat diet-induced obesity. Correspondingly, overexpression of FATP1 in 3T3-L1 cells leads to increased acyl-CoA synthetase activity and FA uptake (Zhan *et al*, 2012), while FATP1 knockdown results in reduced TAG incorporation and a complete loss of insulin stimulated FA uptake (Lobo *et al*, 2007).

Cav1 was identified in a screen of FA-binding proteins (Trigatti *et al*, 1999), and CAV1 and CAVIN1 null mice are lipodystrophic and resistant to high fat diet induced obesity (Liu *et al*, 2008; Razani *et al*, 2002). Similar phenotypes are also detected in humans with mutations in *CAV1* or *CAVIN1* that result in caveola loss. In 3T3-L1 cells, the depletion of Cav1 was shown to result in a 50 % reduction of FA uptake while overexpression resulted in increased FA uptake (Pohl *et al*, 2004; Pohl *et al*, 2002). Furthermore, EHD2, the protein that restricts caveola scission from the PM (Moren *et al*, 2012; Stoeber *et al*, 2012), is upregulated during high fat diet and influences uptake and lipolysis of LCFAs (Matthaeus *et al*, 2020; Moren *et al*, 2019). Since multiple caveola components take part in fatty acid transport, it is tempting to speculate that caveolae *per se* are required for efficient LCFA uptake and metabolism instead of just the individual proteins.

Although the impact of Cav1 and FATP1 on LCFA uptake and metabolism is clear, it is not understood whether their activities are functionally coupled and which precise roles they play in this process. Here, we have addressed the influence and relationship between Cav1 and FATP1 on the early events of LCFA uptake, including PM transport, metabolic trapping and the subsequent processing and effect on lipid synthesis and storage. Using a combination of advanced imaging approaches and quantitative mass spectrometry, we identify and characterize MCSs between caveolae and FATP1-positive ER and their role in LCFA uptake and metabolic processing.

## Results

### FATP1-mediated increase in cellular LCFA uptake and storage is dependent on Cav1

Given the established impact of both Cav1 and FATP1 on the uptake and storage of FAs *in vivo* and *in vitro*, we aimed to determine if their activities were distinct or coupled to the same mechanistic process. We used 3T3-L1 cells, a well-studied pre-adipocyte model that is tractable for exogenous protein expression. Cav1-mRFP and FATP1-Venus were transiently expressed in the cells and their localization was investigated using confocal microscopy. FATP1-Venus displayed a clear ER localization based on the ER marker calnexin (Fig S1A and B), in agreement with previous data (Zhan *et al*., 2012). Cav1-mRFP was mainly localized to caveolae, distinct small puncta on the PM which are also positive for cavin1 (Fig S1A).

To test if increased levels of FATP1 or Cav1 alters the total LCFA uptake in 3T3-L1 cells, Cav1-mRFP or FATP1-Venus-overexpressing cells were incubated with fluorescently labelled palmitic acid (PA-Bodipy) for 1, 5 or 10 minutes, and the total fluorescence intensity per cell was quantified from merged confocal stacks of each cell (Fig 1B and Fig S1C). We observed robust increases in PA-Bodipy uptake in cells expressing either Cav1-mRFP or FATP1-Venus. The increased uptake was evident already after 1 minute of incubation and persisted at 5 and 10 minutes showing that both proteins influence the early stages of FA uptake and metabolic trapping (Fig 1B). The PM localization of Cav1 indicated that Cav1 might be involved in the initial membrane transport. Indeed, confocal microscopy analysis revealed substantial colocalization between PA-Bodipy and Cav1, showing that LCFAs are enriched in caveolae (Fig S1D). Since FATP1-Venus is localized to the ER, it cannot directly manage transport over the PM but rather influences uptake via transport to the ER and / or vectorial metabolic trapping of FA as acyl-CoA. To address if the FATP1-mediated increase in FA uptake was dependent on Cav1, we depleted cells of Cav1 using siRNA and assayed the total uptake with or without FATP1-Venus supplementation. Interestingly, FATP1-Venus expression did not result in increased PA-Bodipy uptake at 1 and 5 min in cells lacking Cav1 (Fig 1B). At 10 minutes, we observed a significant but more modest increase compared to control, suggesting that some LCFA is made accessible to FATP1 via Cav1-independent transport at later timepoints. These data show that the increased early uptake of LCFA mediated by FATP1 is dependent on the presence of Cav1.

To track the fate of internalized FAs in cells, we used palmitoleic acid alkyne (PA-alk) which can be conjugated to the C-171 click-MS reporter (Thiele *et al*, 2019). Thereby, metabolic derivates of PA-alk can be identified and distinguished from endogenous lipids by quantitative mass spectrometry (MS) with signature fragmentation in tandem MS (MS/MS) (Fig 1D). PA-alk was added to 3T3-L1 cells overexpressing Cav1 or FATP1 for 10 minutes, the lipids were extracted, clicked to C-171 and analyzed with MS. The peaks with corresponding fragmentation in MS/MS representing C-171 clicked PA-alk derivatives were identified and their abundance was quantified. We focused the analysis on incorporation into DAGs, TAGs, the phospholipids PC and PI and the sphingolipid SM. PA-alk was incorporated in all these lipid classes (Fig 1D and S2A-E). Strikingly, the overexpression of Cav1 and FATP1 both resulted in 2–3-fold increased incorporation of the PA-alk into TAGs, PCs and PIs compared to control cells (Fig 1D). The level of incorporated PA-alk in all different detected TAGs, PCs and PIs was increased, showing no preference for a specific acyl length (Fig S2B, D, E). We also detected a major increase in double incorporation of PA-alk in TAGs (Fig 1D and S21C) showing that the FA is incorporated into both DAG and TAG. However, the amount of PA-alk in DAG and SM was not increased following Cav1 and FATP1 overexpression (Fig 1D and S2A and C). This suggested that DAG is rapidly metabolized into TAG in wild-type cells and that Cav1 and FATP1 do not promote incorporation into SM. We also investigated if the FATP1-mediated increase was dependent on Cav1 by overexpressing FATP1 in Cav1 KD cells. Interestingly, the increased incorporation in TAG, PC and PI following FATP1 expression was decreased in cells where Cav1 had been depleted (Fig 1D). Instead, the levels of PA-alk in DAGs were increased as compared to controls. This showed that incorporation into DAG is still possible, but that Cav1 is required for the specific FATP1-dependent usage of DAG to generate TAG, PC and PI. For SM we did not detect any significant differences compared to control. Interestingly, Cav1 depletion alone did not result in significant changes in PA-alk incorporation.

Our data suggests that Cav1 and FATP1 work together to facilitate metabolic channeling of DAG to TAG and DAG to PC. As increased TAG levels should lead to more LDs in the cells, we assessed this using fluorescent microscopy. Oleic acid coupled to BSA was added to starved cells and fatty acid incorporation into TAGs after 18 hours was scored by measuring the total LD stain area per image using fluorescent microscopy (Fig 1E-F). Indeed, the LD area was significantly increased in cells overexpressing Cav1 or FATP1. However, FATP1 overexpression was not sufficient to increase LD formation in cells with reduced Cav1 levels, which is in agreement with the MS data. These results suggest that FATP1 and Cav1 act cooperatively to facilitate PM transport, acylation and incorporation of FAs in TAGs and phospholipids.

### FATP1-positive ER is found near caveolae on the PM

To investigate the coupling between Cav1 and FATP1, we overexpressed both proteins and used structure illumination microscopy (SIM), providing 100 nm resolution, to study their localization. FATP1-Venus formed an extensive network consisting of membrane tubules. In the basal plane of cells, these tubules were found to frequently overlap with Cav1-mRFP-positive puncta (Fig 2A). Detailed analysis using 3D SIM revealed that both single caveolae and caveola clusters intersected with the FATP1-positive ER (Fig 2A). This showed that FATP1-positive ER is found in close proximity to the PM and caveolae, but that FATP1-Venus tubules also associated with the PM without Cav1. To determine if the spatial coupling of Cav1 and FATP1 also was linked to the metabolic processing of LCFA into TAGs, we overexpressed FATP1-Venus and stained cells for Cav1 and LDs. Similarly to the Cav1-mRFP, approximately 30% of the endogenous Cav1 fluorescent spot area overlapped with FATP1-Venus ER tubules in the plane of the basal PM (Fig 2B-C). LDs overlapped with FATP1-Venus tubules to a similar extent, which is in line with their ER origin. Yet, the overlap between caveolae and LDs was low in comparison, suggesting that LDs are not found directly at the site of FATP1-caveolae connections (Fig 2B-C). To address potential membrane contact formation in high resolution and in the absence of overexpressed proteins, we performed transmission electron microscopy (TEM) analysis of ultrathin 3T3-L1 cell sections. The ER was indeed detected adjacent to the PM and, strikingly, was surrounded by caveolae, occasionally even in direct contact with them (Figure 2D). We also examined other cultured cell lines similarly and found that in all investigated cell types the ER was in close proximity to caveolae (Fig S3A). Interestingly, we occasionally observed high-density bridge-like connections between the ER and the PM membranes suggesting that proteins might stabilize the MCSs (Figure 2D).

**Figure 2.**
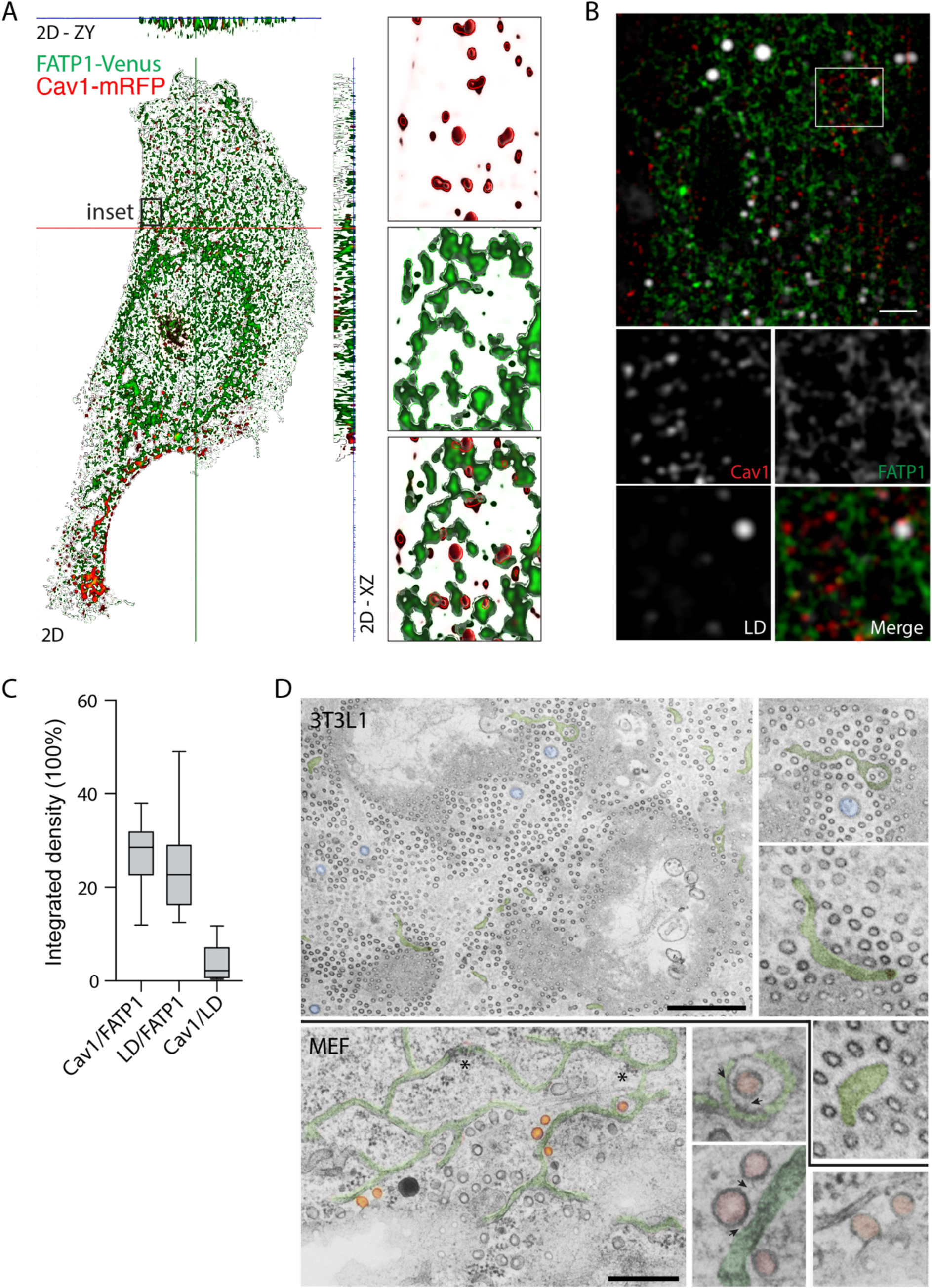
ER positive for FATP1 is found in close proximity to caveolae at the PM. **(A)** Super-resolution lattice SIM image of a 3T3-L1 cell, transfected with FATP1-Venus and Cav1-mRFP, processed for SIM^2^. Whole-cell volume visualized using Zen Black Ortho-slicer, including top view and orthogonal x/y side views. Insets show: (i) Cav1 representing single and clustereded caveolae, (ii) FATP1 network forming a reticular pattern consistent with ER, and (iii) merged overlay showing FATP1 network connecting Cav1 structures. **(B)** Representative SIM image of 3T3-L1 cell expressing FATP1-Venus and stained against Cav1 and LDs (LipidTox). Scale bar, 2 µm. **(C)** Quantification of the co-localization of FATP1 with Cav1, Cav1 with Lipid droplets and FATP1 with Lipid droplets, shown as box-and-whiskers plots. **(D)** Images show glancing sections across the cell surface cut parallel to the cell substratum to illustrate the relationship between ER elements (pseudocolored green) and caveolae (60-70nm spherical uncoated profiles). Upper panels 3T3L1 cell; associated high magnification views of selected regions; Lower panels mouse embryonic fibroblast (MEF); high magnification views of selected regions from main panel on right showing selected caveolae (red highlighting). Black arrows indicate putative bridges between ER and caveolae. Asterisks indicate ribosomes on regions of putative ER tubules. Blue shaded structures in upper panel indicate clathrin coated pits. Scale bars, 1µm upper main figure; 500nm lower main panel.

To further characterize the MCSs between FATP1-positive ER and caveolae we performed correlative light and electron microscopy (CLEM) of 3T3-L1 cells expressing Cav1-mRFP and FATP1-Venus. Cells were unroofed (Fig 3A) and the remaining PM lawns of the basal membrane was prepared for scanning electron microscopy (SEM). The samples were imaged by fluorescence microscopy prior to critical point drying and platinum coating followed by SEM imaging (Fig 3B). Unroofed cells typically displayed a dark circular area with no fluorescence in the middle of the PM lawn as a fingerprint of the previous location of the nucleus (Figure 3B). Cav1-mRFP was detected as distinct puncta suggesting that caveolae were still present in the remaining PM. Furthermore, we observed an FATP1-Venus signal in distinct membranous structures that frequently co-localized with the Cav1-mRFP fluorescence (Figure 3B). This showed that FATP1-positive ER was still associated with the PM following the unroofing process. The fluorescence and SEM images were correlated using semi-automated software (ICY with ecCLEMv2 plugin (Paul-Gilloteaux *et al*, 2017), highlighting areas that were positive for both proteins (Figure 3C). We identified budding clathrin-coated vesicles, clathrin lattices and actin fibers, which confirmed that the PM was relatively intact after the unroofing process (Figure 3C).

**Figure 3.**
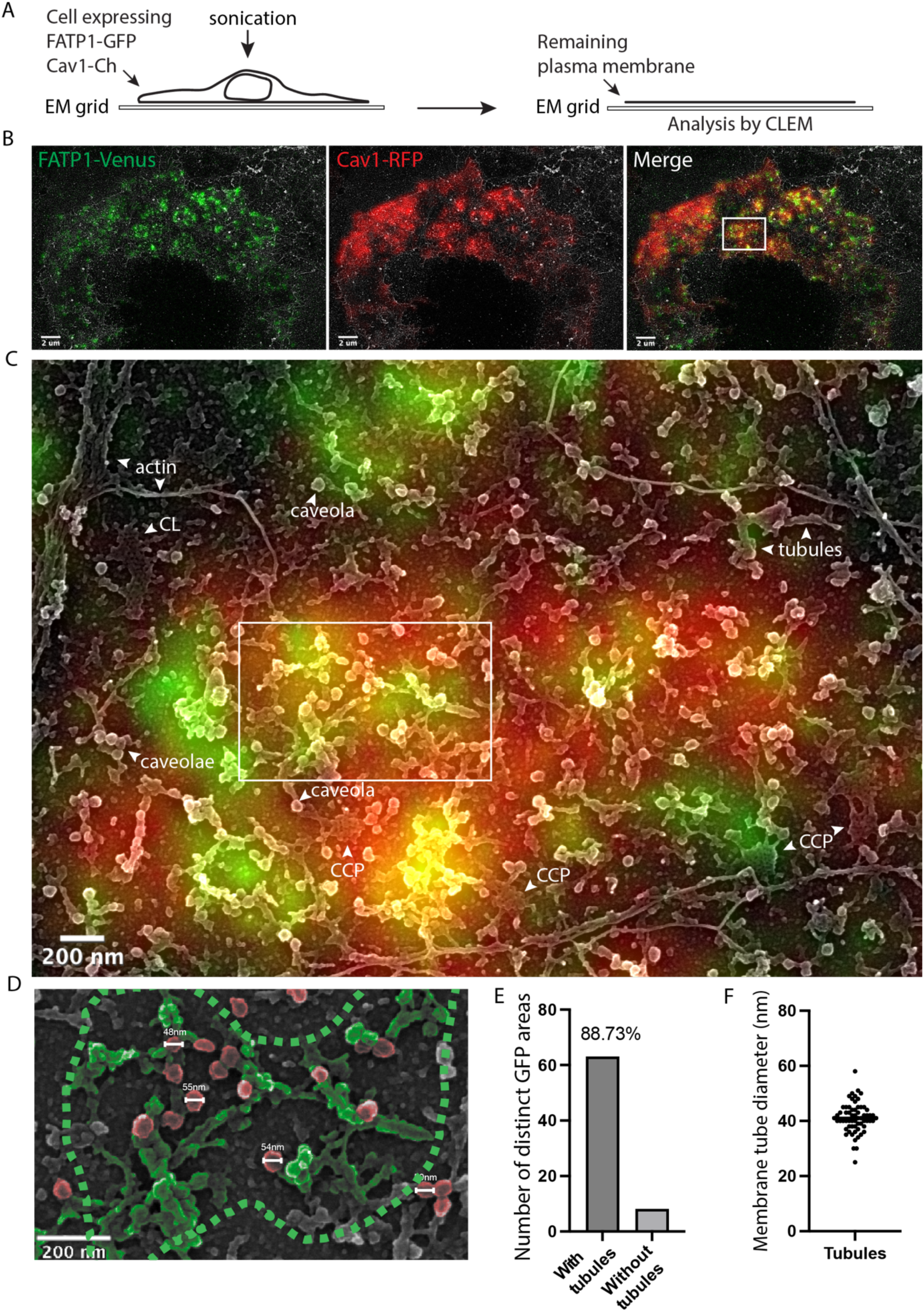
CLEM analysis of PM lawns generated from cells expressing Cav1-mRFP and FATP1-Venus. **(A)** Schematic of the unroofing and correlative light and electron microscopy (CLEM) workflow. **(B)** Correlative SEM and fluorescence images of unroofed cells coated with platinum and imaged after fluorescence acquisition. Upper panels show FATP1, Cav1, and merged fluorescence signals of an unroofed cell (scale bar, 2 µm). **(C)** Inset as indicated by a white square in (B) shows the region at higher magnification. Scale bar, 200 nm. Annotated structural features include actin filaments, clathrin-coated pits (CCP), caveolae, and tubular elements interpreted as residual ER membranes. FATP1-positive regions are enriched in tubular structures, while Cav1-positive regions contain small, round caveolae. Overlapping FATP1 and Cav1 signals correspond to areas with dense networks of interwoven tubules and closely associated caveolae. **(D)** The inset from (C) shows a higher-magnification view with pseudocolored caveolae (red) and ER tubules (green) based on the fluorescence signal. measured caveola diameters are indicated. **(E)** Quantification of FATP1-positive regions in correlative images classified by the presence or absence of tubular structures **(F)** Diameters of FATP1-positive membrane tubules measured across several cells. The average diameter is indicated by a line and individual diameters with dots.

Importantly, the PM was still covered with caveolae (45-55 nm spherical vesicles that correlated with the Cav1-mRFP fluorescence) after the unroofing procedure (Fig 3C and Fig S3C). We focused our analysis on the areas where Cav1 and FATP1 were colocalized according to the fluorescent signals. We detected caveola clusters and ER tubule remnants in these areas of the SEM images (Figure 3D). Quantification showed that 88% of the areas with Cav1 signal (positive for green fluorescence) also contained membrane tubules with diameters of around 40 nm (Fig 3E, F). The tubule size is in agreement with previously reported diameters for peripheral ER tubules (Schwarz & Blower, 2016). Importantly, caveolae were frequently observed in direct contact, or very close proximity with the ER (Figure 3C-D). Taken together, these data suggests that caveolae take part in the formation of MCSs which also harbor the required metabolic enzymes to facilitate efficient FA uptake as well as processing. This model would also resolve the controversy of whether FATP1 localizes to the ER, the PM or both.

### Caveola-ER contact sites are dynamic

To analyze the dynamics of caveola-ER connection sites, we used total internal reflection fluorescence microscopy (TIRFM), which enables rapid live cell imaging of events close (150 nm) to the PM. TIRFM imaging showed almost 30% colocalization between Cav1-mRFP and FATP1-Venus signals at the PM, in agreement with our previous experiments (Fig 4A). We also analyzed ESYT1, an ER protein involved in ER-PM MCS formation (Yu *et al*, 2016). Similar to FATP1, ESYT1-GFP colocalized with Cav1-mRFP (Fig 4B and Fig S4A), which further showed that the ER and caveolae are closely associated. Additionally, we used mitotracker to investigate if caveolae are found near mitochondria, which are key players in FA metabolism. Only few mitochondria were detected by TIRFM and they did not strongly colocalize with the Cav1 signal (Figure 4B and S4B). We tracked the duration and movement of Cav1-mRFP, FATP1-Venus and ESYT1-GFP with time lapse TIRFM using Imaris software. Objects of interest were identified and defined based on their fluorescence intensity and size, with an effective diameter of 400 nm. The center position of each structure in each frame was calculated, and the total movement was visualized by a trace line (1 pixel width and with a voxel size 159 nm) connecting these center positions (Fig 4C). The Cav1 and FATP1 tracks had variable lifetimes with some persisting across the five-minute tracking period (Fig S4C). The tracks frequently overlapped and their mobility seemed to correlate (Figure 4C, Movie 1 and S4C). Interestingly, some FATP1 and caveola tracks appeared to remain close to each other without overlapping during the observation period (Fig 4C).

**Figure 4.**
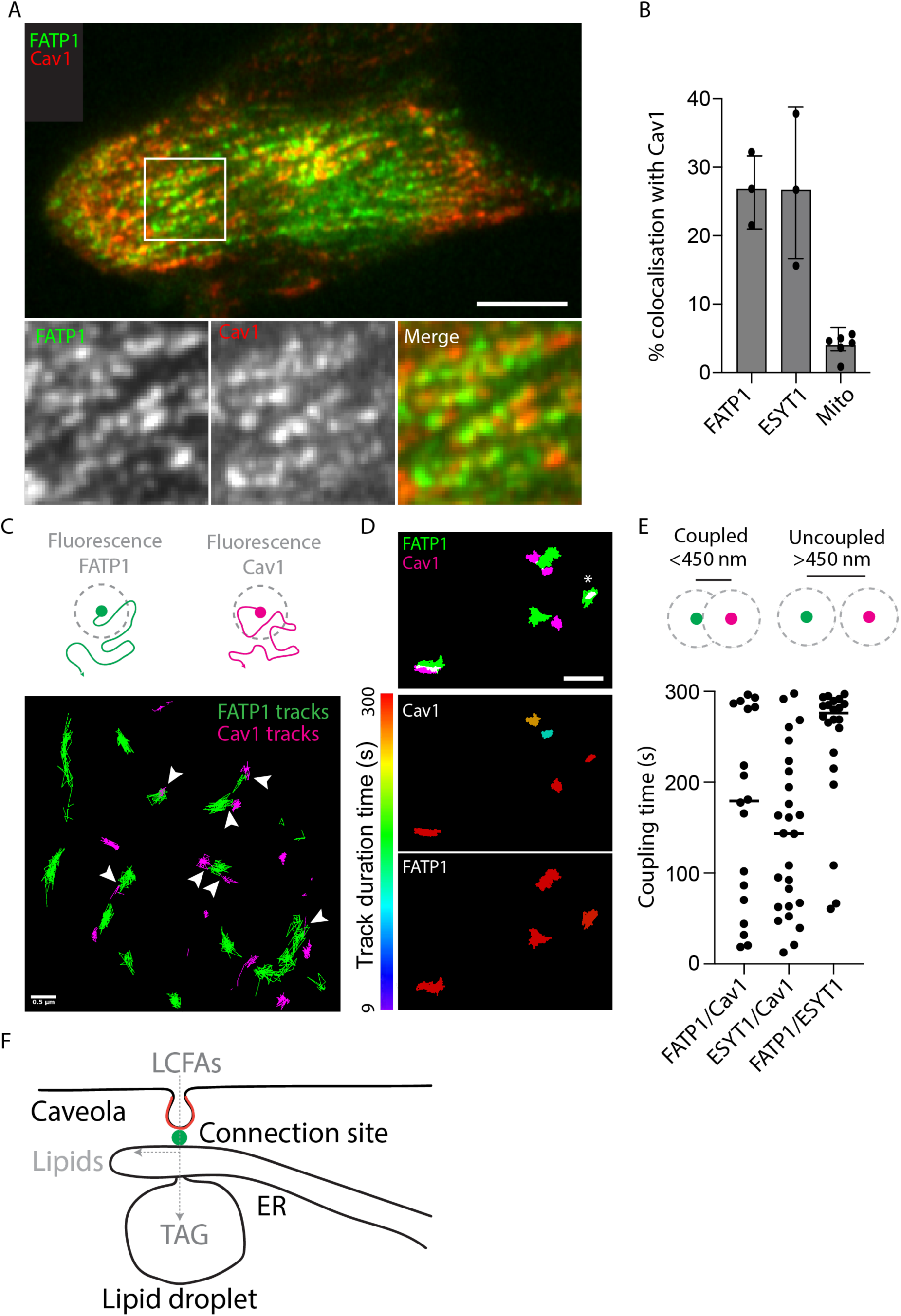
TIRFM analysis reveals dynamic colocalization between Cav1-mRFP and ER associated proteins. **(A)** Micrographs of TIRFM data of Cav1-mRFP and FATP1-Venus co-transfected 3T3-L1 cells. Insets show single channels of FATP1 and Cav1 magnified from indicated area. **(B)** Quantification of the colocalization of Cav1 with FATP1, ESYT1 and MitoTracker in TIRFM. **(C)** Illustration and color-coded center-point analysis of TIRFM data showing tracks from 0-60 seconds from the area indicated in Fig S4C at higher magnification. Arrowheads indicate overlapping. Scale bar, 500 nm. **(D)** Isolated tracks of Cav1 and FATP1 longer than 50 seconds over 5-minute TIRFM movie color-coded by track duration time (top) or merge of isolated tracks of Cav1 (pink) and FATP1 (green). **(E)** Quantification of the coupling time (coupling: centre spots closer than 450 nm as illustrated) between tracks of the indicated proteins as determined by TIRFM and Imaris analysis.

For coupling analysis, we selectively identified tracks longer than 10 s (Fig 4D) and measured the distance between the center positions in each frame and calculated the coupling time based on how long the two center positions remained within a set distance cut-off. As representing each fluorescent signal as a center position leads to an apparent (and erroneous) increase in spatial resolution, we chose a conservative 450 nm coupling cut-off. (Fig 4E). Quantification of the data revealed that most correlated FATP1 and ESYT1 signals persisted for nearly the entire TIRFM movie (mean coupling time 246 ± 73). As both proteins are ER residents (physically linked by the ER membrane), this verifies that the ER is persistently close to the PM. We used their mean coupling time as a cut-off to classify track pairs above this value as strongly coupled. 39 % of FATP1 and Cav1 track pairs formed a distinct subpopulation of strongly coupled signals based on this threshold (Fig 4E). The percent ESYT1 and Cav1 track pairs classified as strongly linked was 27%. Given ESYTs involvement in MCS and based on the strong coupling between ESYT1 and FATP1, it is tempting to speculate that the strongly coupled FATP1/Cav1 track pairs would also contain ESYT1, and that the proteins would work in concert to facilitate MCS formation.

## Discussion

FA molecules are transported almost instantaneously from the PM to the ER during uptake into cells. However, the mechanistic details of how the hydrophobic FAs supposedly cross the aqueous environment between the two membranes remain scarce. In this work, we have investigated an alternative model in which the transport is facilitated by direct PM-ER contacts. Specifically, this work focused on the potential mechanistic coupling of Cav1 (in the PM) and FATP1 (in the ER) to form FA-transporting MCSs.

Using a combination of methods, we measured how Cav1 and FATP1 affect i) the total LCFA uptake at early time points, ii) the metabolic fate of LCFA molecules taken up by the cell, as well as iii) the long-term accumulation of TAGs in lipid droplets. Overall, our results show that the overexpression of either protein leads to increased LCFA uptake, metabolic trapping and storage in TAGs and phospholipids, which is in agreement with previous work (Meshulam *et al*, 2006; Zhan *et al*., 2012). Our data are also in line with the observation that these proteins are overexpressed during adipocyte differentiation and the resulting TAG accumulation in lipid droplets (Pilch *et al*, 2011). While the bulk outcome of overexpressing each protein was similar, the effects varied in time. Raising the cellular FATP1 levels led to a continuous increase in LCFA uptake over ten minutes, whereas Cav1 overexpression had the greatest effect at the one-minute time point. Importantly, the proteins were functionally coupled, as FATP1 supplementation had no effect in the early time points when Cav1 expression was simultaneously reduced. This suggests a directional relationship, where Cav1 initially concentrates the LCFA molecules in caveolae at the PM and subsequently transports them to FATP1 at the ER, especially as we also observed LCFA enrichment in caveolae. Interestingly, the lack of Cav1 was somewhat compensated at the 10 minute timepoint. Slower mechanisms may have been able to take over for Cav1, or the PM may have been saturated with LCFAs due to the high LCFA experimental concentration in the experiment. Furthermore, tracking the metabolic fate of click-labelled LCFA molecules showed that FATP1-mediated LCFA incorporation into TAGs and phospholipids was also affected by Cav1 depletion. In Cav1-deficient cells, LCFA molecules remained trapped as DAGs instead of being processed to TAG (Fig 1A). This shows that Cav1 and FATP1 are not only coupled to facilitate fatty acid uptake but instead their link is critical for the metabolic processing of LCFAs, which is further corroborated by our observation that the FATP1-mediated TAG accumulation in LDs also requires Cav1. Taken together, our data show that Cav1 and FATP1 are intrinsically linked to cooperatively transport fatty acids over the PM, facilitate their acylation, and to ensure their correct incorporation into TAGs at the ER.

The functional link between FATP1 and caveolae begs an obvious question: are they physically connected? Our SIM and TIRFM data show a strong spatial connection between FATP1 and Cav1. The signals were similarly correlated in CLEM, and we could directly observe caveolae in contact with (presumably) ER-derived membrane tubules both in CLEM and TEM. Direct membrane contacts between the endoplasmic reticulum (ER) and the plasma membrane (PM) are thought to play key roles in the transport of membrane lipids (Saheki & De Camilli, 2017). Yet, although it has been proposed that such sites in theory could enable efficient FA uptake (Fullekrug *et al*., 2012), their role in fatty acid metabolism in cells is not known. Our results show that MCSs indeed mediate LCFA transport and indicate that the model could be expanded to downstream LCFA processing instead of just uptake.

The formation, stability and dynamics of PM-ER contact sites remains poorly characterized. Tethering proteins such as ESYT1 have been shown to promote membrane contacts (Saheki & De Camilli, 2017), and while ESYT1 colocalized with Cav1 and FATP1 in our experiments, we cannot conclude if it is required (or sufficient) for MCS formation. We observed (presumably) proteinaceous high-density contacts between caveolae and the ER in EM, although the identity of these connections cannot be determined unambiguously. However, if MCSs are formed by specific protein-protein interactions, the PM and the ER can be strongly coupled. This is indirectly supported by our CLEM observations, as the caveola-ER contacts remained even after the relatively harsh unroofing process. Interestingly, earlier mass spectrometry characterization of purified caveolae is also consistent with strong PM-ER connections (Ortegren *et al*, 2006; Ost *et al*, 2005). The authors described a high-density caveola subpopulation that contained the ER enzymes FATP1 and FATP4, which would have to be tightly bound to remain connected after the rough purification procedure. We observed both short-lived FATP1 and ESYT1-positive puncta showing dynamic ER movements near the PM. However, we also detected signals that remained close to the PM for the entire five-minute observation period. These sites had limited lateral mobility, which suggests that firm connections are formed between the ER and the PM. Additionally, a subpopulation of stable FATP1- and ESYT1-containing puncta remained closely linked to Cav1, showing the presence of caveolae at these sites.

Based on our data, we propose that PM caveolae form direct contacts with the ER to facilitate LCFA transport and metabolism (Fig 4F). First, the LCFAs are enriched in caveolae and efficiently transferred to the ER (without needing to cross an aqueous environment), which is enriched in FATP1, and likely other fatty acid-processing enzymes at these sites. Subsequently, the LCFAs are processed into TAGs and incorporated into LDs. We envision that this enables very efficient and controlled transport and metabolism of LCFAs in adipocytes, where caveolae are highly abundant and the cells are optimized to store TAGs. Yet, we also observed potential MCSs between caveolae and ER MCSs in mouse embryonic fibroblast, primary mouse lymphatic endothelial cells (LEC), endothelia in a capillary of mouse skeletal muscle and human epidermoid carcinoma cells (A431) cells, suggesting that this pathway is not limited to fat cells. However, some cell types such as neurons lack caveolae, indicating that other mechanism and pathways can also facilitate uptake of LCFAs.

Mechanistically, it remains to be addressed if Cav1 and FATP1 are sufficient for the transfer of LCFA between membranes or if this involves other proteins such as FABPs or ACSL1 as well. Also, other fatty acid-metabolizing enzymes may be coordinated to the MCSs. For example, previous results shows that FATP1 and DGAT2 can form complexes (Xu *et al*, 2012). Furthermore, since caveolae endocytosis has also been proposed to facilitate FA uptake (Matthaeus *et al*., 2020), it needs to be clarified if transfer via MCS also would involve scission of caveolae from the cell surface. Interestingly, EHD2, a regulatory caveola protein involved in scission seems to have contradictory roles in lipid transport. EHD2 downregulation leads to increased caveola scission (Moren *et al*., 2012; Stoeber *et al*., 2012) but increased LD size in mice (Matthaeus *et al*., 2020). However, the lack of EHD2 prevents adipocyte differentiation and results in decreased LD size (Moren *et al*., 2019) but leads to epigonadal and peri-inguidal fat accumulation in mice (Matthaeus *et al*., 2020). These apparently conflicting data suggest that caveolae play a regulatory role in the control of both cellular entry and exit of LCFAs. Indeed, transfer mediated by MCSs as we propose, could also favor transport of FA in the opposite way during lipolysis. It is thus tempting to speculate that PM-ER MCSs function both in LCFA import as well as fatty acid export during lipolysis and are therefore highly regulated both on the cellular and the organismal level.

## Acknowledgment

We acknowledge the Biochemical Imaging Center (BICU) at Umeå University within the National Microscopy Infrastructure, NMI (VR-RFI 2016-00968) and the Microscopy Australia Research Facility at the Center for Microscopy and Microanalysis at the University of Queensland for providing support and assistance with microscopy. We thank Irene Martinez at BICU for assistance and expertise with image analysis and data visualization and James Rae (The University of Queensland, Australia) for TEM processing. We thank Maria Ahnlund at the Swedish Metabolomics Centre in Umeå, Sweden for providing support and assistance with mass spectrometry and lipid profiling. We acknowledge the Umeå Core Facility for Electron Microscopy (UCEM), Umeå University and the Swedish National NMR Infrastructure (SwedNMR, 2021-00167). We thank Ho Yi Mak (The Hong Kong University of Science and Technology) for kindly providing the original FATP1-Venus plasmid and Prof. Natasha Harvey for the primary lymphatic endothelial cells. The work was supported by grants to RL from the Swedish Research Council (dnr 2021-05117) the Swedish Cancer Society (CAN 23 3004 Pj 01 H). RGP was supported by an Australian Research Council (ARC) Laureate Fellowship (FL210100107).

## Author contributions

Sebastian Rönfeldt and Richard Lundmark designed the research and Sebastian performed and analysed the cell biology, live cell imaging and tracking experiments and CLEM analysis. Elin Larsson performed and analysed MSMS analysis of LCFA metabolism., Björn Morén performed TIRFM and tracking experiments, Alex Craig synthesized the C171 click reporter and Lindon Moodie supervised, Robert Parton contributed conceptually and enabled the electron microscopy experiments and performed their analysis. Sebastian Rönfeldt and Richard Lundmark conceived the work and wrote the manuscript, and all authors contributed with text, figures and editing.

## Materials and methods

### Cell Lines, constructs and transfections

Murine 3T3-L1 preadipocytes were maintained in Dulbecco’s Modified Eagle Medium (DMEM, high glucose) supplemented with 10% fetal bovine serum (FBS) at 37 °C in a humidified incubator with 5 % CO_2_. Antibiotics were not used in regular culture conditions. For transient overexpression, cells were transfected with plasmid constructs encoding FATP1-Venus, and Cav1-mRFP (all kindly provided by Ho Yi Mak). A FATP1-BFP construct was generated in-house by recloning FATP1 from the FATP1-Venus vector into a BFP backbone. Transfections were performed either by electroporation using the Neon Transfection System (Invitrogen) or with Lipofectamine 3000 (Thermo Fisher Scientific) according to the manufacturer’s instructions. For electroporation, 70-90 % confluent cells were trypsinised and transfected using a 10 µL tip format with the following parameters: 1500 V, 20 ms pulse width, and 2 pulses. The total DNA concentration per electroporation was adjusted depending on the number of constructs: 1000 ng for single, 750 ng per construct for double (total 1500 ng), and 500 ng per construct for triple transfections (total 1500 ng). For Cav1 knockdown, 3T3-L1 cells were seeded in 24-well plates and allowed to adhere overnight. On the following day, cells were transfected with Cav1-targeting siRNA using Lipofectamine 3000. For the first transfection, 50 µL Opti-MEM was mixed with 1 µL of 20 µM siRNA and 1 µL Lipofectamine 3000, incubated for 20 minutes at room temperature, and added to each well. Cells were incubated overnight at 37 °C and 5 % CO₂. Cells were then split into 6-well plates and incubated overnight at 37 °C and 5 % CO₂. A second transfection was performed the next day using 200 µL Opti-MEM with 2 µL of 20 µM siRNA and 2 µL Lipofectamine 3000, incubated for 20 minutes at room temperature, and added to each well. Cells were incubated overnight under the same conditions. Following the second transfection, cells were split and seeded/transfected for subsequent experiments.

### General Chemistry Methods

Thin-layer chromatography (TLC) was performed on 0.2 mm aluminium plates precoated with silica gel 60 F_254_ (Merck). Compounds were visualized with an ultraviolet (UV)-light, and stained with phosphomolybdic acid. Column chromatography was performed with silica gel (40 – 63 μM). ^1^H NMR spectra were recorded at 400 MHz on a Varian Mercury Plus spectrometer. All spectra were recorded from samples in CDCl_3_, at room temperature in 5 mm nuclear magnetic resonance (NMR) tubes. Chemical shifts are reported relative to the residual solvent peak at δ 7.26 for CDCl_3_. Resonances were assigned as follows: chemical shift (multiplicity, number of protons, coupling constant(s)). Multiplicity abbreviations are reported by the conventions: s (singlet), t (triplet), m (multiplet). ^19^F NMR spectra were recorded at 376 MHz on a Varian Mercury Plus spectrometer under the same conditions as for the ^1^H NMR spectra. All solvents and reagents were used as received.

### Synthesis of C171

(Thiele *et al*., 2019)

4-Azido-1-(methylsulfonyloxy)butane

To a stirred solution of 4-azido-1-butanol (0.262 g, 2.59 mmol) in anhydrous CH_2_Cl_2_ (20 mL) was added triethylamine (0.48 mL, 0.35 g, 3.5 mmol) followed by methanesulfonyl chloride (0.20 mL, 0.30 g, 2.62 mmol) under nitrogen at room temperature. The mixture was monitored by TLC, and after 16 hours was diluted into distilled water. The aqueous portion was extracted with CH_2_Cl_2_ (x2), before the organic portions were combined and washed with distilled water (x1) and brine (x1) before the solvent was reduced *in vacuo* to afford the title compound (0.44 g, 83%) as a pale-yellow oil. ^1^H NMR (400 MHz, CDCl_3_) δ 4.27 (t, 2H, *J* = 6.2 Hz), 3.36 (t, 2H, *J* = 6.5 Hz), 3.02 (s, 3H), 1.89-1.82 (m, 2H), 1.76-1.69 (m, 2H).

### C171

A solution of 4-azido-1-(methylsulfonyloxy)butane (0.440 g, 2.96 mmol) and *N*,*N*,*N*-dimethylethylamine (0.31 mL, 0.22 g, 3.0 mmol) in CH_2_Cl_2_ (1 mL) was heated to 60°C under nitrogen, and the mixture was monitored by TLC. After 6 hours, the reaction was cooled and the solvent was reduced in vacuo. The crude mixture was subjected to preparative HPLC (ACN:H_2_O with 0.1% formic acid, 5:95 to 60:40) to afford 0.087 g of the ammonium salt with an undetermined counterion. This material was dissolved in 10 ml 50% MeOH, and loaded onto a 5 ml column of Amberlite A26 (OH^-^-form, primed by washing sequentially with 50% MeOH, 1M NaOH, water, 1M NH_4_BF_4_, water, 50% MeOH) and eluted with 50% MeOH. The first 1.5 ml flow-through were discarded and the following 20 ml collected and reduced *in vacuo* to afford the title compound (0.065 g, 13%) as a yellow oil. ^1^H NMR (400 MHz, CDCl_3_) δ 3.51-3.42 (m, 6H), 3.18 (s, 6H), 1.88-1.80 (m, 2H), 1.72-1.66 (m, 2H), 1.41 (t, 3H, *J* = 7.0 Hz). ^19^F NMR (376 MHz, CDCl_3_) δ −150.8.

### MS sample preparation

3T3-L1 cells were transiently transfected for over expression or depletion of proteins as described above. Cells were plated at 1 x 10^6^ cells per 6-well two hours prior to addition of palmitoleic acid alkyne (Cayman chemicals) at a final concentration of 100 µM in DMEM and chased for 10 minutes. Cells were washed with PBS buffer which was completely aspirated. 500 µl extraction buffer (5:1 v/v methanol:chloroform + internal deuterated standard (ceramide)) was added to the cells which were detached from the wells with a cell scraper. Extracted cells were added to Eppendorf tubes and wells were rinsed with 250 µl extraction buffer (without internal standard) and pooled in designated Eppendorf tube for a final volume of 750 µl. Samples were sonicated for 30 seconds and centrifuged at 20 000 x rcf for five minutes at room temperature. Samples were transferred to new tubes to get rid of cell debris and airdried with N2 gas to a volume of 400 µl. 1 ml of a mix containing chloroform and 1% acetic acid (2:3 v/v) was added to the supernatant followed by 30 seconds of shaking. Samples were centrifuged at 20 000 x rcf for five minutes at room temperature. The nonpolar phase of the samples was transferred to new Eppendorf tubes and dried down at 45°C for 20 min in a SpeedVac (Genevac). Samples were dissolved with 8 µl CHCl_3_ and sonicated for 30 seconds. 40 µl Click-mix (4:1 v/v absolute EtOH: acetonitrile + 20 mM C171, 1 mM Cu(I)TFB) was added to the samples which were sonicated for 30 seconds before a 16 hour incubation at 40 °C. Samples were mixed with 1:1 v/v mix of CHCl3:ddH_2_O and centrifuged at 20 000 x rcf for 5 minutes. Solvent phase was transferred to and dried down in microvials in a SpeedVac for 10 minutes at 45°C and stored at −80°C until analysis.

### LC-MS analysis

The dried samples were resuspended in 100 µl MeOH:CHCl_3_ (3:1 v/v). The chromatographic separation was performed on an Infinity Agilent 1290 (Agilent Technologies, Waldbronn, Germany) ultra-high performance liquid chromatograph coupled with a tandem mass spectrometer (UHPLC-MS-MS). 0.75 μL of each sample were injected onto an Acquity UPLC CSH, 2.1 x 50 mm, 1.7 µm C18 column in combination with a 2.1 mm x 5 mm, 1.7 µm VanGuard precolumn (Waters Corporation, Milford, MA, USA) held at 60 °C. The gradient elution buffers were A (20:80 acetonitrile:water, 10 mM ammonium formate, 0.1% formic acid) and B (89.3:10.5:0.2 2-propanol:acetonitrile:water, 10 mM ammonium formate, 0.1% formic acid), and the flow-rate was 0.5 ml/min. The compounds were eluted with a linear gradient using initial condition 15 % B, and increase to 30 % B at 1.2 min, 55 % at 1.5 min, isocratic to 5.0min, increase to 72 % B at 7 min, 85% at 9.5 min and 100 % B at 10.0 min, and then held at 100 % for 2 minutes. An additional wash of the injection valve, with 100 % B and flow-rate 4.0 ml/min for 70 sec, was performed before decreased to initial condition 15 % B over 0.3 minutes; these conditions were held for 50 seconds to equilibrate the column before next injection. The compounds were detected with an Agilent 6546 Q-TOF mass spectrometer equipped with a jet stream electrospray ion source operating in positive ion mode. The exact masses of individual lipid molecules were detected with an Agilient 6550 Q-TOF mass spectrometer equipped with an iFunnel jet stream electrospray ion source (Agilent Technologies). A reference interface was connected for accurate mass measurements; the reference ions purine (4 μM) and HP-0921 (Hexakis (1H, 1H, 3H-tetrafluoropropoxy) phosphazine) (1 μM) were infused directly into the MS at a flow rate of 0.05 ml/min for internal calibration, and the monitored ions were purine m/z 121.05 and HP-0921 m/z 922.0098. The first batch of extracts was analyzed in positive mode and then in negative mode before second batch was injected. The flow gas temperature was set at 150°C, the drying gas flow to 8 l/min and the nebulizer pressure to 35 psig. The sheath gas temp was set to 350°C and the sheath gas flow 11 l/min. The capillary voltage was set to 4000 V in positive ion mode. The nozzle voltage was 300 V. The fragmentor voltage was 120 V, the skimmer 65 V and the OCT 1 RF Vpp 750 V. The collision energy was set to 25 and 40 V. The m/z range was 100 - 1700, and data was collected in centroid mode with an acquisition rate of 4 scans/sec. All data processing and peak identification was performed using the Agilent Masshunter Profinder version B.10.0.2 (Agilent Technologies Inc., Santa Clara, CA, USA). A LC-MS library of exact masses and experimental retention times of peaks corresponding to the expected masses of labeled lipid classes combined with the characteristic neutral loss was built up and used for identification. The extracted features were aligned and matched between samples with a m/z tolerance of 20 ppm and a retention time window of 0.01 min. Results were expressed as area under the curve (ACU) values from the extracted ion chromatograms of each lipid molecule normalized to the ACU of the internal standard within the same sample. Statistical analyses were performed using GraphPad Prism, T-test, unpaired parametric test.

### Fluorescence microscopy and immunostaining

Fluorescence imaging was performed using either a Zeiss spinning disk confocal microscope equipped with a 63× oil immersion objective, or a Leica SP8 Falcon confocal laser scanning microscope with a 63× oil immersion objective. Excitation and emission settings were selected according to the respective fluorophores. Images were acquired using identical microscope settings for all comparative conditions. Acquired images were processed and analyzed using Fiji (ImageJ) for visualization. Structured Illumination Microscopy (SIM) was performed on a Zeiss Elyra 7 microscope operating in lattice SIM mode with 13-15 phases and a Plan-Apochromat 63×/1.4 Oil DIC M27 objective. Excitation was achieved using 488 nm and 561 nm laser lines for Venus and mRFP detection, respectively. Post-processing was performed in ZEN Black software using SIM² reconstruction, and ZEN Blue was used for 3D rendering and orthogonal slicing. For Colocalization analyses cells, Images were acquired as whole cell Z-stacks using a Zeiss Elyra 7 in lattice SIM mode with a Plan-Apochromat 63×/1.4 Oil DIC M27 objective. Laser lines used were 405 nm, 488 nm and 642 nm for the respective stains, followed by SIM processing in Zen Black. Final images were search for the basal layer Z-slice and thresholded for Cav1, FATP1 and LipidTox signals using ImageJ. Thresholded Signals were overlaid and reduced to only overlapping regions for Cav1/FATP1, Cav1/LipidTox and FATP1/LipidTox combinations. Integrated density of overlapping signals was calculated and plotted as percentage values.

For immunostaining, cells were fixed with 4 % paraformaldehyde (PFA) for 15 min at room temperature, followed by three washes with phosphate-buffered saline (PBS). Cells were then permeabilized, washed and incubated in either 0.1% Triton X-100 in PBS for 10 min (a-calnexin) or 0,05 % saponin in PBS for 20 min (a-Cav1) in all subsequent steps. Blocking was done in 5 % goat serum for 1 h at room temperature and incubation with the primary antibodies rabbit anti-Calnexin (Sigma-Aldrich, dilution 1:200), or rabbit anti-Cav1 (Abcam, dilution 1:500) was done for 1 h at room temperature. Following three washes, cells were incubated for 1 h with Alexa Fluor 647-conjugated anti-rabbit secondary antibodies (Invitrogen, dilution 1:1000). Lipid droplets were labeled using HCS LipidTOX Deep Red Neutral Lipid Stain (LipidTox) (ThermoFisher Scientific, dilution 1:1000) for 1 h. Coverslips were then washed three times, rinsed once with Milli-Q water and mounted on glass slides using Dako Fluorescence Mounting Medium (Agilent Technologies) and stored at 4 °C until imaging.

### Analysis of PA-Bodipy uptake and lipid droplet formation

For analysis of PA-Bodipy uptake, 3T3-L1 cells cultured on glass coverslips were transiently transfected by electroporation 21 hours prior to analysis by electroporation as described above. Following starvation for 2 hours in serum-free DMEM, cells were incubated with 10 µM C16-Bodipy (ThermoFisher Scientific) (dissolved in medium containing BSA) for 1, 5, or 10 minutes at 37 °C. Subsequently, cells were washed with phosphate-buffered saline (PBS) and fixed with 4 % paraformaldehyde (PFA) for 20 minutes at room temperature. Fixed cells were washed three times with PBS, mounted on slides, and imaged using a spinning disk confocal microscope using a 63× objective with 488 nm laser for Bodipy, 405 nm for BFP, and 568 nm for mRFP detection. Whole-cell z-stacks were collected, and maximum intensity projections were generated using Fiji. Transfected cells were segmented using Cellpose, and mean fluorescence intensity was measured with Fiji. Values were normalized to control, and statistical analyses were performed using GraphPad Prism (one-way Anova).

For analysis of lipid droplet formation, 3T3-L1 cells were transfected using Lipofectamine 3000 and incubated overnight at 37 °C and 5% CO₂. Prior to lipid loading, cells were starved for 2 hours in serum-free medium and subsequently incubated with ∼0.3 mM oleic acid (Sigma-Aldrich) complexed to BSA for 18 hours to induce lipid droplet formation. Cells were then fixed and stained with BODIPY 493/503 (Invitrogen) to visualize lipid droplets. Fluorescence images were acquired using a 40× objective under identical exposure settings across conditions in three independent experiments generating datasets suitable for quantitative comparison. Image analysis was performed using Fiji (ImageJ). Images were background-corrected and thresholded to segment lipid droplets, and the total lipid droplet area per image was quantified. Measurements were based on the entire field of view without segmentation of individual cells or consideration of transfection status. Statistical analyses were performed using GraphPad Prism (one-way ANOVA).

### TIRF live-cell imaging and tracking analysis

Cells were transferred to live-cell imaging chambers with coverslip bottoms and maintained in phenol-red-free live-cell medium at 37 °C during acquisition. TIRF live-cell imaging was performed using an inverted Nikon Eclipse Ti-E inverted microscope with an DU897 ANDOR EMCCD camera controlled by Nikon NIS Elements interface and equipped with 488 nm and 561 nm laser lines for excitation of Venus and mRFP, respectively, perfect focus system and a 63× oil-immersion objective. Time-lapse image sequences of the basal membrane of cells were recorded continuously over 5 minutes at 1 s intervals. Image post-processing and quantitative tracking analysis were carried out using the Imaris software 10.0.1 tracking module (Bitplane). Structures with a diameter of 0.4 μm were tracked and the applied algorithm was based on Brownian motion with max distance traveled of 0.45 μm and a max gap size of 3. Tracks in figures were displayed using different colormaps as detailed in the figure legends. For track-coupling analysis, tracks within smaller regions of interest (ROIs) were manually verified and curated for missing track connections at 1px gauss to ensure continuous tracking and accurate motion analysis. Tracks shorter than 10 s in each ROI were discarded. The spatial distance between individual track spots from two different imaging channels in each time frame were compared using Imaris software. Particle trajectories that remained within a radius of 450 nm were classified as coupled and the coupling time was measured. The data included 5 different ROIs from two different cells per condition. Graphical representation was performed using GraphPad Prism.

### Correlative light and scanning electron microscopy (CLEM) and transmission electron microscopy

3T3-L1 cells were co-transfected with FATP1-Venus and Cav1-mRFP using electroporation transfection and seeded on glass coverslips pretreated with 0.01 % poly-L-lysine for 30 min and dried overnight prior to transfection. Cells were cultivated overnight at 37 °C in a humidified incubator with 5 % CO₂ and the following day, cells were washed with HEPES-buffered saline (HBS) containing protease inhibitors (without EDTA) and briefly incubated with 0.01 % poly-L-lysine for 20 s. Coverslips were rinsed with Buffer (2mM CaCl_2_, 1mM MgCl_2_ in HBS) for 30–60 s, hypotonic buffer for 30 s and HBS with protease inhibitors. Cells were unroofed by sonication using a sonotrode (1 s, 20 % amplitude) and subsequently washed in HBS containing protease inhibitors. Cells were fixed with 4 % PFA for 30 min at room temperature and washed three times with HBS containing protease inhibitors. Fluorescence imaging was performed in cell chambers containing HBS and protease inhibitors using a spinning disc confocal microscope. After imaging, cells were washed with Milli-Q water and prepared for scanning electron microscopy (SEM). Coverslips were dried using a Leica EM CPD300 critical point dryer, mounted onto SEM stubs using silver glue, and coated with a 2 nm platinum layer using a Quorum Q150T-ES sputter coater. SEM imaging was performed on a Carl Zeiss Merlin FESEM. For correlative image analysis, fluorescent and SEM images were aligned using Icy software with the ec-CLEM v2 plugin. Areas of interest were annotated and color-coded based on morphological features and fluorescence signals using GIMP. CLEM image analysis was carried out manually using multiple imaged regions obtained from different cells. FATP1-Venus fluorescence was first used to identify regions of interest, after which the diameters of the corresponding structures were measured directly in the electron microscopy channel. To further validate the nature of these FATP1-enriched regions, fluorescence signal areas were examined and classified based on whether associated tubule structures were present or absent in the EM images. This approach enabled the distinction between FATP1-positive tubules and other fluorescent structures within the field of view. All image inspection, measurements, and classifications were performed using the ICY bioimage analysis platform.

For conventional transmission electron microscopy, cells or tissues were fixed in 2.5% glutaraldehyde in PBS and then processed and imaged as described previously [PMID: 32709891]. Primary mouse lymphatic endothelial cells were prepared as described previously (Betterman JCI 2020 PMID: 32182215), kindly provided by Professor Natasha Harvey (University of South Australia and SA Pathology, Adelaide, Australia). Cultured cells were processed for electron microscopy in 3cm dishes *in situ* and sections were prepared parallel to the substratum.

